# Role of NS1 antibodies in the pathogenesis of acute dengue infection

**DOI:** 10.1101/348342

**Authors:** Deshni Jayathilaka, Laksiri Gomes, Chandima Jeewandara, Geethal S.Bandara Jayarathna, Dhanushka Herath, Pathum Asela Perera, Samitha Fernando, Ananda Wijewickrama, Clare S. Hardman, Graham S. Ogg, Gathsaurie Neelika Malavige

**Affiliations:** Centre for Dengue Research, University of Sri Jayewardenepura, Nugegoda, Sri lanka; National Institute of Infectious Diseases, Angoda, Sri Lanka; Weatherall Institute of Molecular Medicine, John Radcliffe Hospital, Oxford, OX3 9DS, UK

## Abstract

The role of NS1-specific antibodies in the pathogenesis of dengue virus infection is of particular interest to the dengue field, yet remains poorly understood. We therefore investigated the immunoglobulin responses of patients with dengue fever (DF) and dengue hemorrhagic fever (DHF) to NS1. Antibody responses to recombinant-NS1 were assessed in serum samples obtained throughout illness of patients with acute secondary DENV1 and DENV2 infection by ELISA. NS1 antibody titres were significantly higher in patients with DHF compared to those with DF for both serotypes, during the critical phase of illness. Antibody responses were further assessed to NS1 peptides and showed that in both acute secondary DENV1 and DENV2 infection, the antibody repertoire of DF and DHF patients is directed towards distinct regions of the NS1 protein. Further experiments in healthy individuals, with either past severe dengue or past asymptomatic dengue infection revealed that individuals with past inapparent disease mounted antibody responses directed to the same NS1 epitope regions as those with mild acute infection (DF). Our results suggest that the specific epitope target of NS1-antibodies generated by patients could predict disease severity and be of potential therapeutic benefit in aiding vaccine and treatment design.

## Introduction

Dengue virus (DENV) is one of the most rapidly emerging viral infections worldwide, infecting 390 million individuals annually. Despite such obvious need, there is currently no licensed specific drug for treatment of this potentially fatal disease ^1^. A tetravalent live attenuated yellow fever and dengue chimeric vaccine has been recently licensed in some countries, however it has poor efficacy in naïve individuals and efficacy depends on DENV serotype ^2^. Indeed, the vaccine manufacturer recently suggested that this vaccine should be avoided in dengue naïve individuals, due to the likelihood of disease enhancement ^3^. One of the major challenges faced in the development of an efficacious dengue vaccine is our poor understanding of what constitutes a protective immune response.

DENV infections result in a spectrum of disease ranging from subclinical inapparent presentation, through mild dengue fever (DF) to severe dengue hemorrhagic fever (DHF), which is characterized by an increase in vascular permeability resulting in shock and organ dysfunction ^4^. Understanding the molecular pathway that leads to development of vascular leak would provide a major step forward in the development of effective dengue treatments. One such potential target is dengue non-structural protein 1 (NS1), a secreted dengue glycoprotein that is used as a diagnostic marker of disease appearing early in the serum before antibodies are generated^5^. NS1, synthesized as a monomer, forms a dimer in the ER lumen and hexamer in the serum. NS1 is thought to function as a cofactor for viral RNA replication^6^. The mechanism by which NS1 contributes to disease is not fully established and remains controversial. NS1 has been shown to trigger cytokine release and contributes directly to vascular leak through binding TLR4 and engaging the endothelial glycocalyx ^7,8^. In addition, *in vitro* data show that some antibodies against NS1 cross-react with endothelial cells and induces apoptosis, which is suggested to contribute to endothelial dysfunction and vascular leak ^9^. Interestingly, NS1 is important for viral evasion of complement activation by binding mannose-binding lectin and preventing neutralization of DENV via the lectin pathway ^10^. However, some studies have shown that NS1 activates complement, a process that is further enhanced by NS1-specific antibodies ^11^.

Secondary dengue infections incur an increased risk of developing DHF, which supports the hypothesis that NS1-specific antibodies derived from primary infection may, upon expansion following secondary challenge, play a role in disease pathogenesis ^12,13^. However, apoptosis of the endothelium and deposition of immune complexes in the endothelium have not been demonstrated in autopsy studies of fatal dengue ^14^. In contrast to the data in the above studies, some investigations in dengue mouse models have shown that mice injected with sera from dengue infected mice or monoclonal antibodies to NS1, have significantly reduced vascular leak, implying a protective role for antibodies to NS1 ^15^. It is currently not clear if antibodies targeted to specific regions of dengue NS1 protein may play differential roles in the protection or pathogenesis of dengue fever in humans.

The majority of human DENV-specific antibodies are directed against DENV envelope and PrM proteins, and these responses have been relatively well characterized^16–19^. However, NS1 specific antibodies have not been thoroughly investigated despite constituting 27% of antibody responses ^20^. There are limited data on the target, isotype and function of anti-NS1 antibodies in acute secondary dengue infections, which account for most cases of severe dengue infection.

Interestingly, Dengvaxia®, the only registered dengue vaccine currently available, does not generate DENV-NS1-specific antibodies ^21^. and some have speculated that the failure to raise NS1-specific antibodies may contribute to the lower than expected efficacy of this vaccine ^22^. In order to address these questions, here we investigate NS1-specific antibody responses in a cohort of patients with acute secondary dengue (DENV1 and DENV2) infection and study how NS1-specific antibody responses evolve during the course of illness in relation to clinical disease severity. We further characterize anti-NS1 antibody responses in DF and DHF patients and in healthy individuals with varying severity of past dengue infection to identify epitope regions within the NS1 protein sequence that associate with disease progression or protection.

## Results

### Dengue patient cohort

To investigate NS1 antibody responses in patients with acute secondary dengue infection we recruited 76 patients with acute dengue, and through analysis of patient IgM: IgG antibody ratios we determined that 55/76 patients had a secondary dengue infection (IgM: IgG antibody ratio <1.2). These individuals were included in our analyses of NS1 antibody responses. 20 of the secondary dengue infection cohort had acute DENV-1 infection and 35 had acute DENV-2 infection. 10 patients with acute DENV1 infection had DHF, while 10 had DF and 23 patients with an acute DENV2 infection had DHF and 12 had DF (supplementary table 1).

### Kinetics of NS1 antibody production in patients with varying severity of acute secondary DENV1 and DENV2 infection

Serum samples were collected daily from the patients within our cohort, and levels of DENV1 (Fig. 1a) or DENV2 (Fig. 1b) NS1-specific IgG antibodies were measured. We found that NS1 antibody responses in both acute secondary DENV1 and DENV2 infection increased profoundly during the critical phase and were significantly higher throughout in DHF patients compared to those with DF (Fig. 1a and Fig. 1b). In DENV1-infected DHF patients the serum concentration of NS1 antibodies started to rise 5.5 days after the onset of illness. NS1 antibody levels were significantly higher in patients with DHF than in those with DF on day 6 (p=0.003) and 7 (p=0.002) of illness, which represents the critical phase (Fig. 1a). The same pattern was observed in patients with an acute secondary DENV-2 infection, although the difference in antibody titres between patients with DHF and DF were less significant than in DENV1 infection (day 6 p=0.04). It has been shown previously that onset of the critical phase in patients with DENV-2 infection occurs significantly earlier than in acute DENV1 infection ^23^. Accordingly, here we find that NS1 antibody titres rise a day earlier in DHF patients with DENV-2 (day 4.5) than DENV1 infection (Fig. 1b and 1a).

**Figure 1:**
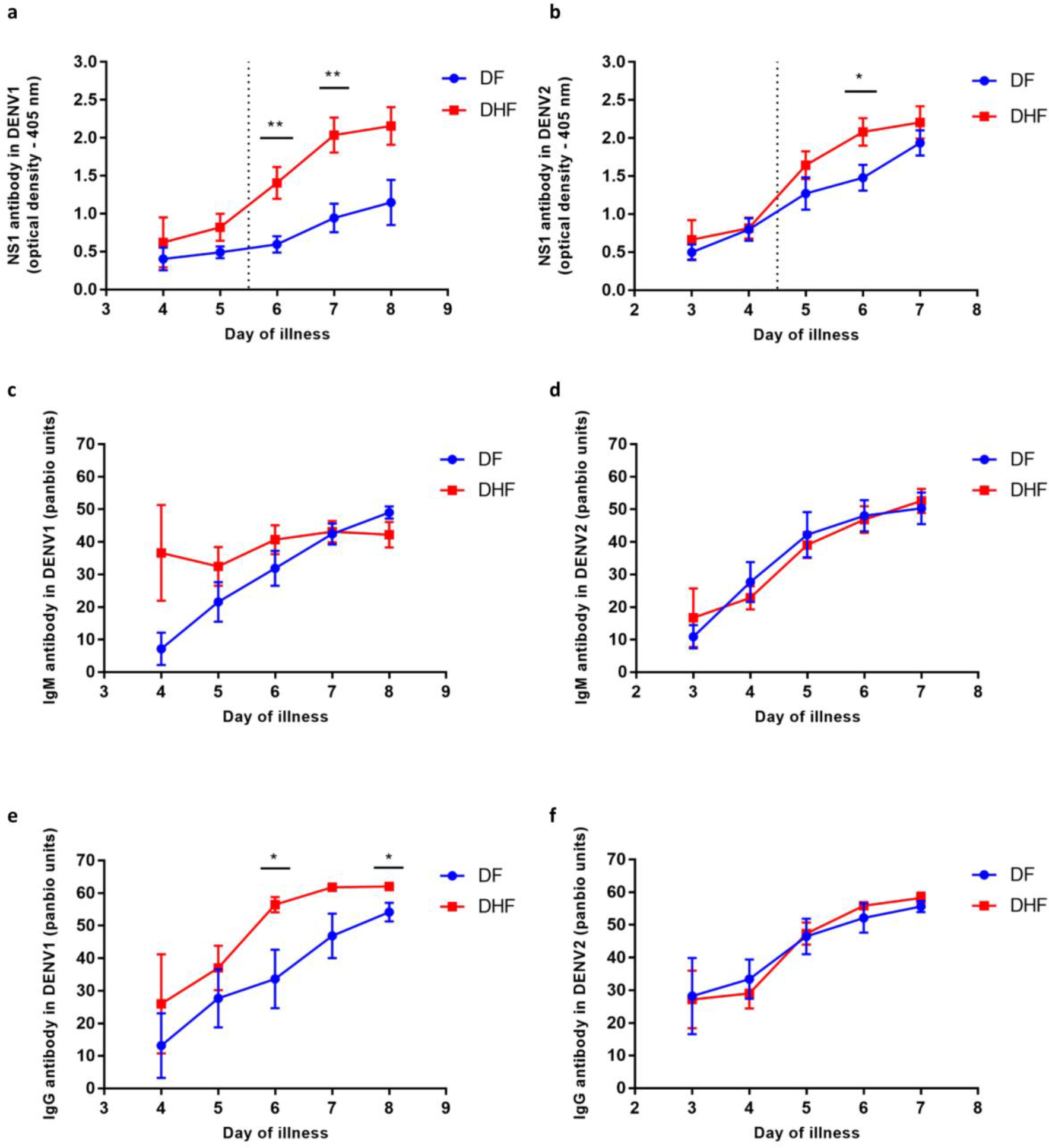
Kinetics of DENV NS1 specific antibody responses in patients throughout the course of illness with varying severity of acute secondary DENV1 and DENV2 infection. (A) DENV1 NS1-specific IgG antibody levels, measured in patients with an acute secondary DENV1 infection causing DF (n=10) or DHF (n=10) using ELISA. (B) DENV2 NS1-specific IgG antibodies levels were measured in patients with an acute secondary DENV2 infection causing DF (n=12) or DHF (n=23) using ELISA. DENV specific IgM antibody (Panbio unit) in patients with an acute secondary DENV1 (C) and DENV2 (D). DENV specific IgG antibody levels were measured in patients with an acute DENV1 (E) or DENV2 (F) infection. Differences between serial values of NS1 specific antibodies in patients with DHF and DF were compared using the Holm-Sidak method. The vertical dotted line represents the day in which the patients entered the critical phase. NS1 antibody levels in patients with DHF are indicated in red, and in those with DF in blue. Error bars indicate mean and standard error of mean (SEM). *P<0.05, **P<0.01

We also semi-quantitatively measured DENV-specific IgM and IgG antibody levels throughout the course of illness, using the PanBio IgM and IgG capture ELISA, in the patients with acute secondary DENV2 infection. Both Panbio IgM and IgG ELISAs use DENV envelope protein as the coating antigen and antibody titres are expressed as PanBio units. We detected no significant difference in DENV-specific IgM antibody titres throughout the course of illness between patients with DHF or DF with either acute secondary DENV1 or DENV2 infection (Fig. 1c and 1d respectively). Interestingly, in patients with an acute secondary DENV1 infection, IgG antibody titres were significantly higher during the critical phase on day 6 (p=0.019) and day 8 (p=0.018) (Fig. 1e). However, we detected no difference in the DENV-specific IgG antibody titres in patients with an acute secondary DENV2 infection (Fig. 1f).

NS1 antibody titres of DHF and DF patients with acute secondary DENV2 infection inversely correlated with platelet counts (Spearman’s r=−0.3, p=0.008 and Spearman’s r=−0.3, p=0.03 respectively) (supplementary fig 1a and 1b). In acute secondary DENV1 infection, NS1 antibody titres inversely correlated with the platelet counts in patients with DHF (Spearmans r=−0.3, p=0.02), but there was no significant correlation in patients with DF (Spearmans r=−0.1, p=0.54) (Supplementary fig 1c and 1d).

### Characterization of antibody responses to NS1 peptides in patients with acute dengue

Our observation that NS1 antibody responses were significantly increased in patients with acute secondary DENV1 and DENV2 infection during the critical phase of illness, lead us to investigate the epitopes recognized by anti-NS1 antibodies. To characterize the antibody targets, serum samples were taken on day 7 of illness from patients with DENV1 infection, and day 6 of illness for DENV2 infection, to align with the critical phase of illness. This was where the most significant difference in NS1 antibody titres was observed between DF and DHF patients. We performed ELISA on patient serum samples to assess antibody recognition of a series of overlapping peptides constituting the entire length of the NS1 protein sequence.

We found that in both acute secondary DENV1 and DENV2 infection, we detected significantly higher antibody titres in DF patients (Fig. 2 and 3) recognizing certain regions, or peptides, of NS1 compared to DHF patients. The epitope regions, for which a significant difference in antibody responses between DF and DHF patients was detected, were similar for DENV1 and DENV2 overlapping NS1 peptides (Fig. 2 and Fig. 3). Patients with DF had significantly higher responses to DENV-1 NS1 peptides spanning a.a 73-107 (pep14, 15 and 17) and a.a 115-137 (pep 21-22) regions and DENV-2 NS1 peptides a.a 33-111 (pep 5-13), a.a 115-141 (pep 16 and pep 18) and a.a 146-163 (pep 21) (Fig. 2b, 2c, and 3a-3c). In addition, patients with DF had significantly higher antibody titres to the epitope region represented by a.a 152-180 of the both DENV-1 and DENV-2 NS1 (peptide 26-31 in DENV-1 NS1 and peptides 21-24 in DENV-2 NS1) (Fig. 2d and 3c).

**Figure 2:**
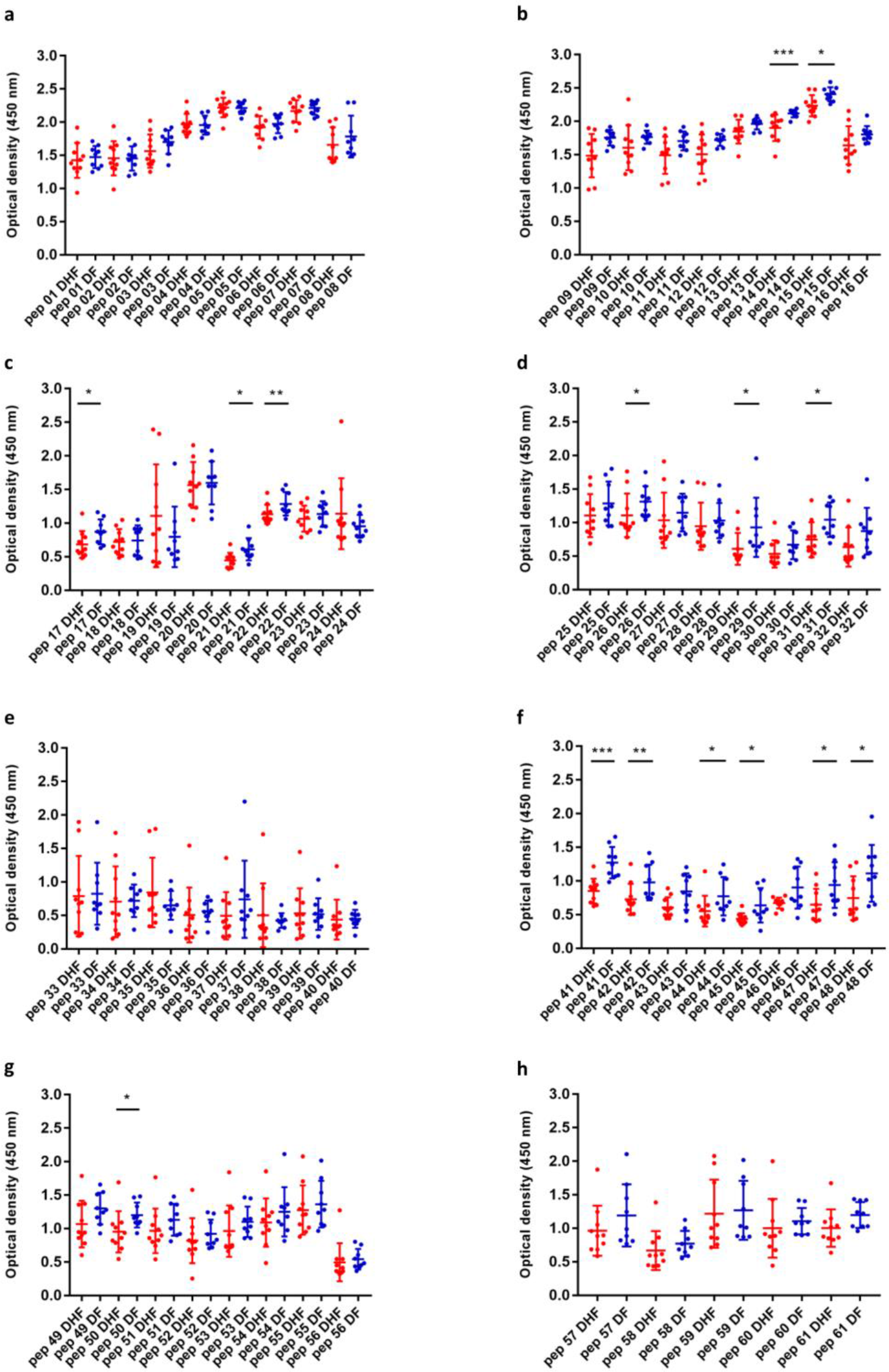
Characterization of DENV1 NS1-specific IgG antibody responses to DENV1 NS1 overlapping peptides in patients with acute secondary DENV1 infection. (A-H) DENV1 NS1-specific IgG antibody responses to 61 DENV-1 NS1 overlapping peptides in patients with DF (n=10) and DHF (n=10) were measured by ELISA on day 7 of illness. Differences in mean values between antibody responses to overlapping peptides in patients with DF and DHF were compared using the Mann-Whitney U test (two tailed). NS1 antibody levels in patients with DHF are indicated in red, and in those with DF in blue. Error bars indicate mean and standard deviation (SD). *P<0.05, **P<0.01, ***P<0.001

**Figure 3:**
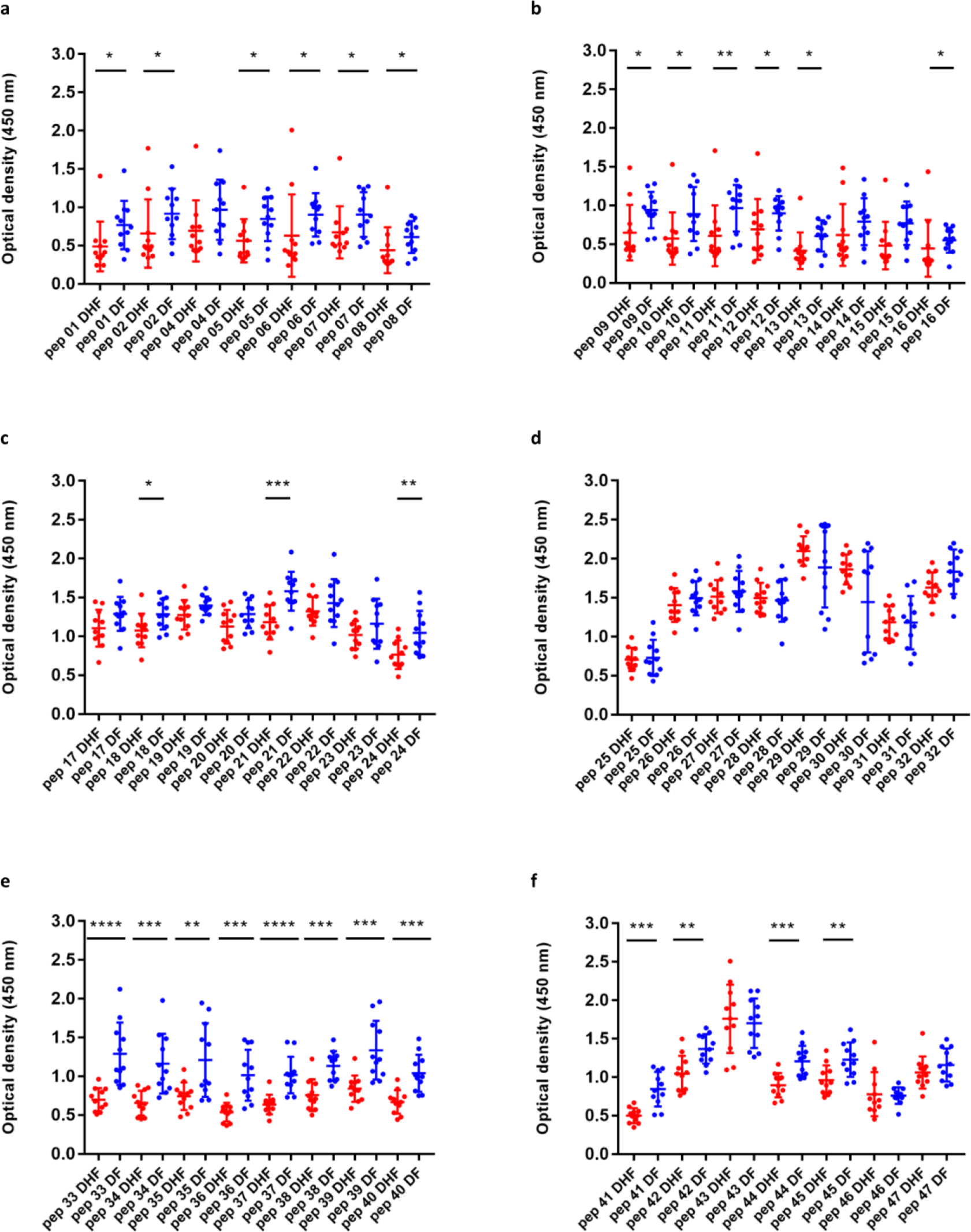
Characterization of DENV2 NS1-specific IgG antibody responses to DENV2 NS1 overlapping peptides in patients with acute secondary DENV2 infection. (A-F) DENV2 NS1-specific IgG antibody responses to 47 DENV-2 NS1 overlapping peptides were measured by ELISA in patients with DF (n=10) and DHF (n=10) on day 6 of illness. Differences in mean values between antibody responses to overlapping peptides in patients with DF and DHF were compared using the Mann-Whitney U test (two tailed). Error bars indicate the means and standard deviation (SD). *P<0.05, **P<0.01, ***P<0.001

Notably patients with DF, as a result of both DENV1 and DENV2 infection, had significantly higher antibody titres to the distal C-terminal region (a.a 230-300) represented by pep 41-48 of DENV-1 and pep 33-42 and pep 44-45 in DENV-2 (Fig. 2f and 3e). In contrast, no differences were observed between patients with DF and DHF with DENV1 or DENV2 to the proximal end of C terminal NS1 (Fig. 2e and 3d).

### Kinetics of NS1 antibody responses to NS1 peptides throughout the course of acute dengue

Having observed that patients with acute secondary DF, compared to DHF, generated more antibodies targeted toward similar epitope regions of DENV1 and DENV2 protein sequences during the critical phase of illness, we sought to investigate how the antibody responses to these regions evolve through the course of illness, by evaluating the responses to these protein sequences before the critical phase (febrile phase), during the critical phase and then the recovery phase. Therefore, we chose 7 regions of NS1 in DENV1 and DENV2, represented by a combination of 2 or 4 peptides to detect antibody specificity, for which patients with DF made significantly higher antibody responses. We aimed to see the kinetics of the antibody responses to these particular regions, to find out if the higher responses to these regions by patients with DF, was limited to the critical phase, or if it was associated with recovery. In both DF and DHF patients, responses to pool 2 (aa 121-142) followed by pool 7 (aa 341-353) elicited the highest antibody titre reading (Supplementary figure 2). Patients with DF had higher responses to peptide pools 2 (0.02) and pool 7 (p=0.03), representing the C terminus end of NS1, towards the end of the critical phase than DHF sera, suggesting that antibody responses to these immunodominant regions of NS1 could associate with protection.

### Determining if NS1 antibody epitopes were predominantly conformational

Although we found that antibodies to recombinant NS1 were significantly higher in patients with DHF during the critical phase, further analysis of NS1 epitopes showed that patients with DF, had significant higher antibody titres to certain peptides/regions of NS1. In order to determine whether the NS1 antibodies bind to conformational epitopes not represented in the peptide binding assay, , we coated 96 well plates with recombinant or denatured DENV NS1 and measured the antibody titres in the sera of patients with DF and DHF during acute secondary DENV1 infection. The recombinant NS1 was denatured by heating the protein at 95° C for 15 minutes as previously described^24^. The sera used for these experiments were sampled on day 7 of illness from patients with DENV1 and day 6 from patients with DENV2. The detection of antibody titres significantly decreased when the DENV-1 NS1 protein capture antigen was denatured in patients with acute DENV1 infection resulting in DHF (p= <0.0001) and DF (p=0.008) (Fig. 4a). Similarly, a significant reduction in the antibody titres was seen when DENV2-NS1 protein was denatured in patients with DHF (p=0.0004). However, no significant difference was observed in those with DF (p=0.17), although a similar trend was observed (Fig. 4b).

**Figure 4:**
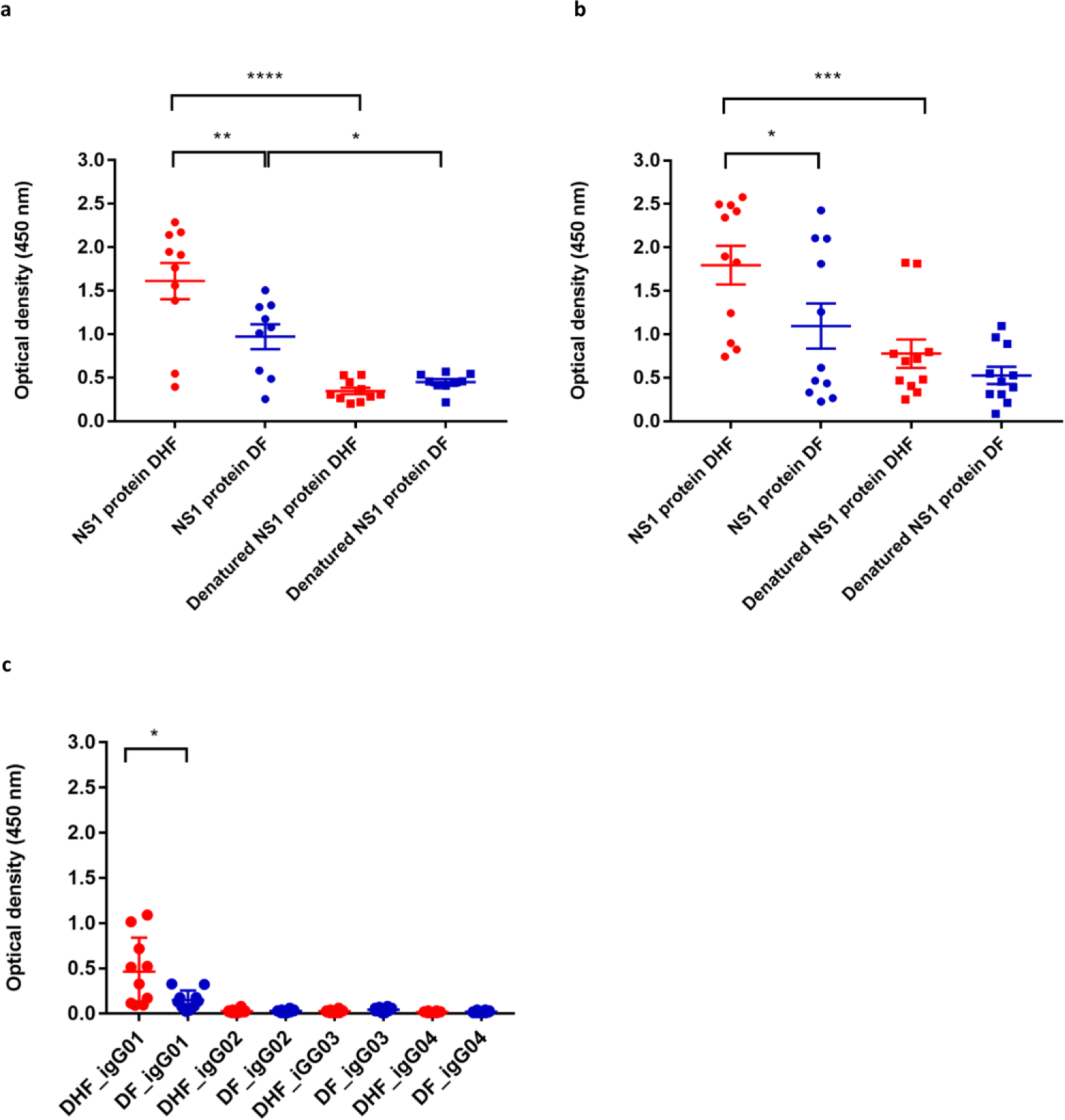
DENV NS1 antibody responses to recombinant DENV NS1 and denatured DENV NS1 in patients with acute secondary dengue and identification of antibody subclass. (A) DENV 1 NS1-specific IgG antibody responses to recombinant DENV 1 NS1 and denatured DENV1 NS1 were measured by ELISA in patients with DF (n=10) and DHF (n=10) on day 7 of illness. (B) DENV2 NS1-specific IgG antibody responses to recombinant DENV2 NS1 and denatured DENV2 NS1 were measured by ELISA in patients with DF (n=10) and DHF (n=10) on day 6 of illness. (C) DENV2 NS1-specific IgG subclasses to DENV-2 NS1 protein were measured in patients with DF (n=10) and DHF (n=10) on day 6 of illness. Differences in mean values between antibody responses to NS1 protein in patients with DF and DHF were compared using the Mann-Whitney U test (two tailed). Error bars indicate mean and standard deviation (SD). *P<0.05, **P<0.01, ***P<0.001.

### Determining NS1 IgG subclass types in patients with acute dengue

Different IgG subclasses that form immune complexes with the antigens, vary in their ability to activate complement and have also been shown to have different affinities for the various IgG receptors ^25^. While IgG binding to FcγRs such as FcγRI, FcγRIIA/C, FcγRIIIB results in activation of macrophages, monocytes and dendritic cells, IgG binding to FcγRIIB inhibits immune responses ^26,27^. Among the IgG subclasses, IgG3 has shown to be the predominant subclass that activated the inhibitory FcγRIIb receptors, whereas IgG1 preferentially activated the activating receptors ^28^. Since our results showed that NS1 IgG antibodies of patients with DHF significantly increased during acute secondary dengue, which coincided with the critical phase, we proceeded to investigate the specific IgG subclasses in patients with acute secondary DENV2 infection by identifying the NS1 specific IgG antibody bound to the microwells by using different biotinylated antibody specific to different subclasses. We found that IgG1 was the predominant subclass of NS1 antibody present and NS1-specific IgG1 was significantly higher in patients with DHF (p=0.03), when compared to patients with DF (Fig. 4c). In contrast, the NS1 IgG3 subclass levels showed a trend to be higher in patients with DF (median 0.04, IQR 0.3 to 0.6) compared to those with DHF (median 0.02, IQR 0.01 to 0.03) although this was not significant (p=0.1). We detected no differences in the other IgG subclasses between patients with DF and DHF.

### Characterization of antibody responses to NS1 overlapping peptides in healthy individuals with varying severity of past dengue infection

Recurrent dengue infection is thought to be a risk factor for developing severe forms of dengue fever. In addition, the presence of antibodies to a previously infecting viral serotype are thought to exacerbate disease upon reinfection. We characterized the antibody responses in healthy individuals, 20 of whom were dengue seronegative individuals, 24 dengue seropositive who had experienced an inapparent dengue infection (NSD) and 17 individuals who had recovered from an episode of DHF (SD). We cannot know when the NSD cohort contracted DENV or whether the infection was primary or secondary, due to their lack of clinical presentation.

As observed in patients with acute dengue infection (Fig 2 and 3), we found that healthy participants with past SD and NSD recognized distinct regions of DENV-NS1. We observed negligible levels of antibodies to the NS1 overlapping peptides in DENV-seronegative individuals (Fig. 5a to o). Patients with past SD, had significantly higher antibody responses to regions represented by peptides 1 to 4, 9 to 16, 22 to 26 and 29 to 31. In contrast, as observed in patients with milder forms of acute infection (DF), those who had past NSD had significantly higher antibody responses to the distal C terminus of NS1 compared to those with past SD, represented by peptide 33 to 37, 40 to 43, 45 to 48 and 58 to 60 (Fig 5 a to o). Therefore, while those with past SD, had significantly higher antibody responses to the N-terminus of the NS1 antigen those with past NSD, had significantly higher antibodies to the peptides representing the C-terminus (supplementary figure 3).

**Figure 5:**
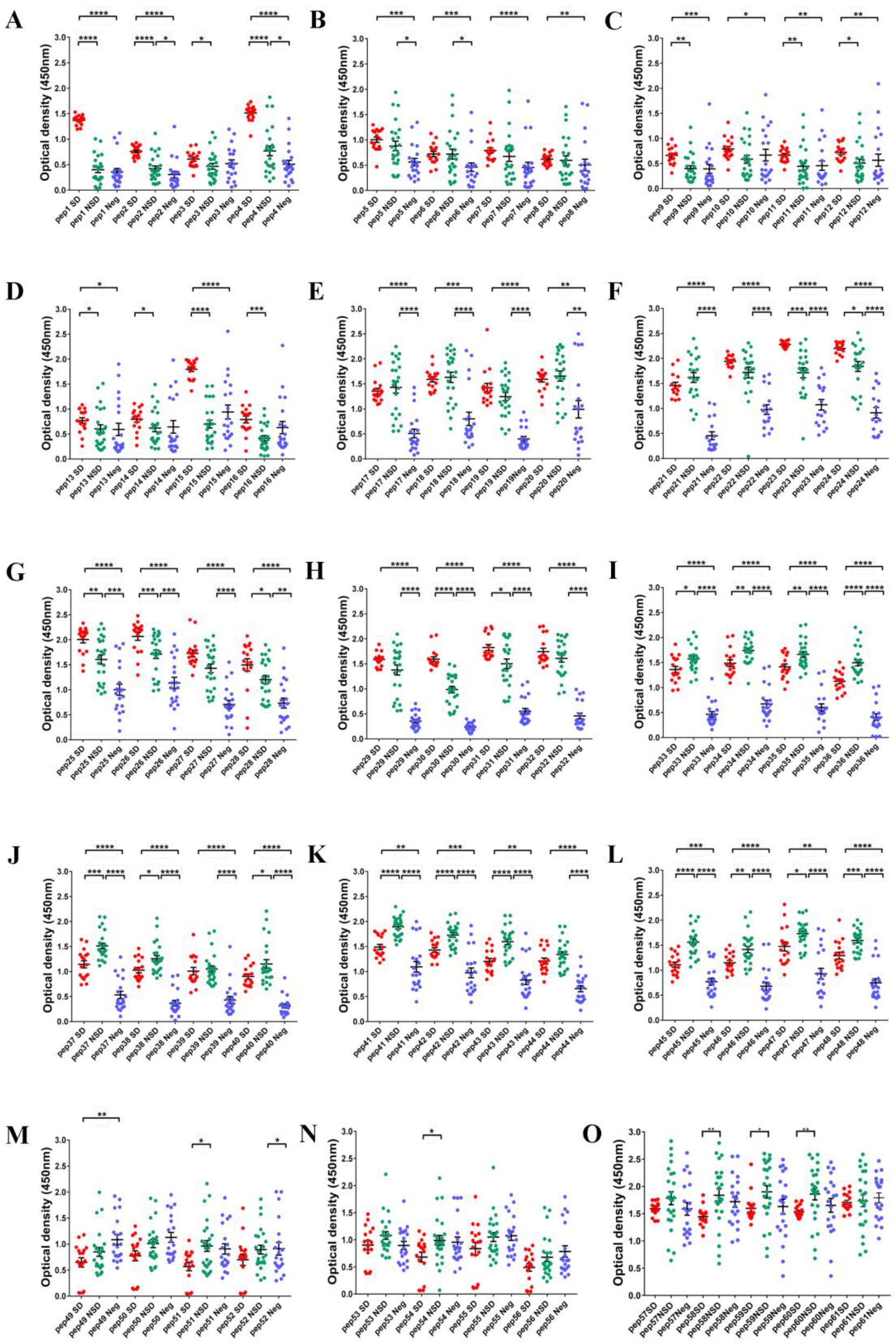
Characterization of DENV1 NS1-specific IgG antibody responses to DENV1 NS1 overlapping peptides in individuals with past infection (A-O). DENV1 NS1-specific IgG antibody responses to DENV-1 NS1 overlapping peptides 1-32 in seronegative individuals (Neg, n=20), those with past non-severe dengue (NSD, n=24) and past severe dengue (SD, n=17). Error bars represent mean and standard deviation. Differences between the groups were compared using the Mann-Whitney U test (two tailed). *P<0.05, **P<0.01, ***P<0.001, ****P<0.0001

## Discussion

In this study we sought to improve our understanding of the specificity of NS1 antibody responses in mild and severe dengue infection disease states. We found that NS1 antibody levels rose more quickly and to a greater extent in patients with DHF during acute secondary dengue infection in the critical phase of illness compared to those with DF. Interestingly, disease severity was reflected in the epitope bias of the antibody response, patients with DF and DHF displayed antibody responses to distinct regions of DENV-NS1 protein, as defined by the constitutive peptide fractions. When we further characterized NS1 antibody responses in a cohort of healthy individuals who either had past DHF or past inapparent infection (NSD), we found similarly that participants with past SD and NSD mounted antibody responses that recognized different regions of NS1, when measured ex vivo to NS1 peptides. Interestingly, those with milder forms of acute dengue and past NSD generated antibody responses to shared regions of NS1, distinct to the epitopes recognized by antibody responses of those with current or past severe acute dengue. This suggests that antibody recognition of certain epitope regions of dengue NS1 protein could confer a greater protective response to dengue infection.

Many of the studies that show NS1 antibodies are associated with protection have been carried out in dengue mouse models ^15,29^. Proteomics, sequence identity analyses and in vitro studies have shown that the C-terminal region of NS1 contains many cross-reactive epitopes, with sequence homology to self-antigens such as endothelial cell proteins and mediators involved in the coagulation pathways and platelet response ^9,30–32^. As a result, the monoclonal antibody, which substitutes the C-terminus of dengue NS1 with that of Japanese Encephalitis NS1 was found to be protective in dengue mouse models ^33^. Our data show that DF patients with acute secondary DENV-1 and DENV-2 infection, had significantly higher antibody titres to the C-terminus of NS1 compared to those with severe forms of dengue (DHF). In fact, when we studied the antibody response kinetics to the C-terminus in patients with DHF and DF, we observed that the levels of antibody directed at the C-terminus of NS1, rose significantly preceding the recovery phase of DF patient illness, whereas DHF responses had declined by this time point. In addition, healthy participants who experienced past NSD also had significantly higher antibody titres directed towards the C-terminus of NS1. Therefore, although the C-terminus appears to share cross-reactive regions with self-proteins, milder forms of illness associates with higher antibody titres targeted to this region. However, it is noteworthy that in our experiments, we used NS1 overlapping peptides which might not reflect the actual binding of NS1 antibodies to the fully folded and polymeric structure. This indeed may account for our observation that antibody responses to the whole recombinant NS1 protein are greater in DHF, whereas we detected higher antibody responses to NS1 peptides in serum of patients with DF. Therefore, in order to further characterize the possible protective and pathogenic antibodies to NS1, it would be important to map antibody epitopes to the dimeric and hexameric forms of NS1.

Although we observed similarities in the NS1 antibody responses from patients with acute secondary DENV1 and DENV2, we did observe some interesting differences. For instance, the difference in NS1 antibody titres during the critical phase between patients with DF and DHF was more marked during DENV1 infection. In addition, while those with DF, due to either serotype, had higher antibody titres to the C-terminus region of NS1, DHF patients with DENV2 infection had significantly higher antibody responses directed towards the N-terminus. Although all DENV serotypes are known to cause severe disease, the severity of infection and clinical features have been shown to vary depending on the viral serotype and genotype ^23,34,35^. During this study period, it was found that patients with DENV2 had a higher risk of developing DHF, and more rapid progression to plasma leakage ^23^. Therefore, the targets of NS1 antibodies generated to each serotype, may play a role in disease pathogenesis.

It has been shown that NS1 antibodies bind to and result in destruction of platelets in dengue mouse models ^36,37^. Our data showed a trend towards inverse correlation between NS1 antibody levels and platelet counts in patients with DHF. Although the possibility that NS1 antibodies lead to thrombocytopenia in humans has not been investigated, one possible explanation for our observations is that NS1 antibodies could be binding to platelets in patients with DHF, causing their destruction. Since the NS1 antibody repertoire appears to be different in those with DF compared to DHF, it is possible that patients with DHF may have NS1 antibodies that preferentially bind platelets. Therefore, careful experiments should be carried out to determine whether such pathways may contribute to disease pathogenesis.

Dengue mouse models have suggested that NS1 antibodies were associated with protection, yet human in vitro experiments have cast some doubt on this hypothesis. Studies of human sera from those with mild and severe forms of dengue, have shown that the binding of NS1 antibodies to endothelial monolayers was significantly enhanced when using sera from patients with severe dengue during the critical phase of illness, a possible route to inducing life-threatening plasma leak ^9^. Furthermore, NS1-NS1 antibody complexes have been shown to activate complement ^11^. and these complexes are known to cross-react with platelets, which could be linked to the hemorrhage seen in severe disease. Therefore, it has been speculated that NS1-antibody complexes are involved in disease pathogenesis ^32,38^. Our data show that NS1 antibodies become elevated in patients with DHF earlier than DF. DHF patient NS1 antibody titres rise as they enter the critical plasma leakage phase and remain significantly higher than in patients with DF, who do not develop vascular leakage. Since the preferential NS1 antibody binding epitopes are different in patients with DHF and DF (and in those with past SD and NSD), it is possible that patients with DHF develop antibodies to certain regions of the NS1, which are associated with enhanced complement activation and endothelial dysfunction, whereas patients with DF produce NS1 antibodies directed against regions of NS1 that neutralize the pathogenic action of NS1. Therefore, it would be crucial to study monoclonal antibodies generated in patients with varying severity of illness during acute infection and assess the ability of these antibodies to activate complement or to neutralize the pathogenic effects of NS1.

In summary, our data show that NS1 antibody titres rise in patients with acute secondary dengue with the onset of the critical phase of disease. Notably anti-NS1 antibody production rises more quickly and to a greater extent in patients with the most severe dengue infection. In addition, antibodies from patients with more severe forms of acute illness and those who have had past DHF recognize different epitope regions of NS1 protein compared to those from patients who have or have previously experienced milder forms of acute dengue. These data suggest that patients with more severe forms of dengue could have a qualitatively and quantitatively different anti-NS1 antibody repertoire, which contributes to disease enhancement. These findings have implications for approaches to treatment and vaccination.

## Methods

### Patients

Adult patients with varying severity of acute dengue infection were recruited from the National Institute of Infectious Disease, Sri Lanka following informed written consent. Daily consecutive blood samples were obtained from 76 patients from the day of admission to the hospital until they were discharge from the hospital, capturing the course of illness. The day on which the patient first developed fever was considered day one of illness. Only those whose duration from onset of illness was ≤4 days were recruited. All clinical features, including presence of fever, abdominal pain, vomiting, bleeding manifestations, hepatomegaly, blood pressure, pulse pressure and evidence of fluid leakage, were recorded several times each day. Fluid leakage was assessed by ultra sound scans to detect fluid in pleural and peritoneal cavities. Clinical disease severity was classified according to the 2011 World Health Organization (WHO) dengue diagnostic criteria ^39^. Patients with a rise in haematocrit > 20% of baseline or clinical or ultrasound scan evidence of plasma leakage were classified as having DHF. Shock was defined as having cold clammy skin, along with a narrowing of pulse pressure of 20 mmHg. As such, 36 patients were classified as DHF and 40 patients were classified as DF.

### Healthy individuals with varying severity of past dengue

In our previous studies, we had recruited 1689 healthy individuals attending the Family Practice Centre, which is a primary health care facility of the University of Sri Jayewardenepura, Sri Lanka, providing community health care to over 2000 families living in the suburban areas of the Colombo district ^40^. In this cohort we have serum samples from the time of recruitment to the study, information regarding dengue serostatus, and whether the individuals had DHF or mild or asymptomatic dengue. Twenty serum samples were used from those who reported an episode of DHF (past severe dengue (SD)) and 24 serum samples were used from those who were seropositive for dengue, but had never been hospitalized due to a febrile illness and (past inapparent dengue infection (NSD)). We also used 20 serum samples from individuals who were seronegative at the time of recruitment.

### Ethics statement

Ethical approval was obtained by the Ethics Review Committee of the Faculty of Medical Sciences, University of Sri Jayawardenapura. All patients were recruited following informed written consent.

### Serotyping of DENV and assessment of viral titre

Acute dengue infection was confirmed by quantitative real time PCR and DENV viruses were serotyped and titres quantified as previously described ^41^. Briefly, RNA was extracted from serum samples using QIAamp Viral RNA Mini Kit (Qiagen, USA) according to the manufacturer’s protocol. Multiplex quantitative real-time PCR was performed as previously described using the CDC real time PCR assay for detection of the dengue virus ^42^, and modified to quantify DENV. Oligonucleotide primers and a dual labeled probe for DEN 1,2,3,4 serotypes were used (Life technologies, India) based on published sequences ^42^. In order to quantify viruses, standard curves of DENV serotypes were generated as previously described in Fernando, S. *et.al* ^41^.

### Analysis of dengue NS1 antigen and DENV IgM and IgG levels

The presence of NS1 antigen was assessed using the NS1 early dengue enzyme-linked immunosorbent assay (ELISA) (Panbio, Brisbane, QLD, Australia). NS1 antigen levels were semi quantitatively assessed in serial blood samples of patients and expressed as PanBio units. Dengue antibody assays were performed using a commercial capture-IgM and IgG ELISA (Panbio, Brisbane, Australia) ^43,44^. Based on the WHO criteria, patients with an IgM: IgG ratio of >1.2 were considered to have a primary dengue infection, while patients with IgM: IgG ratios <1.2 were categorized under secondary dengue infection ^45^. The DENV-specific IgM and IgG ELISA was also used to semi-quantitatively determine the DENV-specific IgM and IgG titres, which were expressed as PanBio units.

### Development of an ELISA to detect NS1 specific antibody

Currently there are no assays to measure NS1 antibody levels, as such we developed an ELISA to measure NS1 antibody levels in patient samples. Commercially available recombinant DENV-1 and DENV-2 NS1 full length recombinant proteins (Native antigen, USA) expressed from a mammalian cell line was used to coat the ELISA plates and capture NS1 specific antibodies in serum samples. The optimal concentration of coating antigen, serum dilution and concentration of secondary antibody were determined by checkerboard titrations.

Our indirect ELISA to detect dengue NS1 specific antibodies was performed in 96 well microtiter plates (Pierce™ Cat: 15031). The plates were coated with either 100μl/well DENV-1 NS1 protein (Native antigen, USA) or DENV-2 NS1 protein (Native antigen, USA) diluted in carbonate-bicarbonate coating buffer (pH 9.6) at a final concentration of 1μg/ml and incubated overnight at 4°C. After incubation the wells were washed three times with 300μl/well washing buffer (phosphate buffered saline (PBS) containing 0.05% Tween 20). The wells were blocked with 300μl/well with blocking buffer (PBS containing 0.05% Tween 20 and 1 % Bovine serum albumin (BSA) for 1 hour at room temperature. The wells were washed again three times with 300μl/well washing buffer. Serum samples were diluted 1:5000 in PBS containing 0.05% Tween 20 and 1 % BSA. Diluted serum samples were added to the appropriate plate at 100μl/well and incubated for 60 min at room temperature.After incubation, wells were washed three times with 300μl/well washing buffer. Goat anti-human IgG, biotinylated antibody (Mabtech, Sweden) was diluted to 1:1000 in PBS containing 0.05% Tween 20 and 1 % BSA and added 100μl/well. The plates were incubated again for 30 min at room temperature and washed 3 times with washing buffer. Streptavidin Alkaline Phosphatase (Abcam, UK) was diluted 1:20 with PBS containing 0.05% Tween 20 and 1 % BSA and added 100μl/well. The plate was incubated for 30 min at room temperature and washed 5 times. Para-nitro-phenyl-phosphatase (PNPP) (Thermo Fisher Scientific, USA) substrate was added at 100μl/well and incubated for 20 mins in dark at room temperature. The reaction was stopped by adding 50μl/well 2M NaOH, and the ELISA read on MPSCREEN MR-96A ELISA reader.

### Detection of antibody responses to DENV1 and DENV2 NS1 overlapping peptides

Dengue NS1 peptide arrays were obtained through the NIH Biodefense and Emerging Infections Research Resources Repository, NIAID (NIH: Peptide Array, Dengue Virus Type 1, Singapore/S275/1990, NS1 Protein, NR-2751 and NIH: Peptide Array, Dengue Virus Type 2, New Guinea C (NGC), NS1 Protein, NR-508). Ninety-six-well microtiter plates (Pierce™ Cat:15031) were coated with 100μl/well peptide preparations diluted in bicarbonate/carbonate coating buffer (pH 9.6) at a final concentration of 1μg/100μl. The peptides were coated individually and incubated overnight at 4°C. After incubation, unbound peptide was washed away twice with phosphate-buffered saline (PBS, pH 7.4). The wells were blocked with 250 μl/well PBS containing 0.05% Tween 20 and 1% Bovine serum albumin (BSA) and incubated for 1.5 hrs at room temperature. The plates were then washed three times with 300 μl/well washing buffer (PBS containing 0.05% Tween 20). Serum samples were diluted 1/250 in ELISA diluent (Mabtech, Cat: 3652-D2). Diluted serum samples were added 100 μl/well and incubated for 30 min at room temperature. After incubation, wells were washed three times with 300 μl/well washing buffer. Goat anti-human IgG, biotinylated antibody (Mabtech, Cat: 3820-4-250) was diluted 1/1000 in PBS containing 1% BSA and added 100 μl/well. Plates were then incubated for 30 min at room temperature and washed three times as described previously. Streptavidin-HRP (Mabtech, Cat: 3310-9) was diluted 1/1000 in PBS containing 0.1% BSA and added 100 μl/well. Incubation was carried out for 30 min at room temperature and washing procedure was repeated five times. TMB ELISA substrate solution (Mabtech, Cat: 3652-F10) was added 100 μl/well and incubated in the dark for 10 min at room temperature. The reaction was stopped by adding 100 μl/well 2M H_2_SO_4_ stop solution and absorbance values were read at 450nm.

### Analysis of antibody binding to NS1 epitopes

DENV1 NS1 and DENV2 NS1 recombinant proteins were added into separate wells in carbonate-bicarbonate coating buffer (pH 9.6) at a final concentration of 1μg/ml and was incubated overnight at 4°C. NS1 proteins, which were denatured by heating them for 95°C for 15 minutes as previously described^24^, were used to compare the responses to the recombinant NS1 in this assay. The wells were blocked with 300μl blocking buffer (PBS containing 1 % Bovine serum albumin (BSA)) for 1 hour at room temperature. DENV1 and DENV2 serum samples diluted 1:5000 with ELISA diluent (Mabtech, Cat: 3652-D2). Diluted serum samples were added at 100 μl/well into respective wells and incubated for 30 min at room temperature. The bound NS1 specific antibodies were detected using 100 μl/well Goat anti-human IgG, biotinylated antibody (Mabtech, Sweden), diluted 1:1000 in PBS containing 1 % BSA, and incubated for 30 min at room temperature. Streptavidin-HRP (Mabtech, Cat: 3310-9) diluted 1/1000 in PBS containing 0.1% BSA was added 100 μl/well and incubated for 30 min at room temperature. TMB ELISA substrate solution (Mabtech, Cat: 3652-F10) was added 100 μl/well and incubated in the dark for 10 min at room temperature. The reaction was stopped by adding 100μl/well 2M H_2_SO_4_ stop solution and absorbance values were read at 450nm.

### Characterization of DENV2 NS1-specific IgG subtype antibody responses

DENV2 NS1 recombinant protein was bound to microplate wells with carbonate-bicarbonate coating buffer (pH 9.6) at a final concentration of 1μg/ml and was incubated overnight at 4°C. The wells were blocked with 300μl blocking buffer (PBS containing 1 % Bovine serum albumin (BSA)) for 1 hour at room temperature. DENV2 serum samples diluted 1:5000 in ELISA diluent (Mabtech, Cat: 3652-D2) were added 100 μl/well and incubated for 30 min at room temperature. The bound NS1 specific IgG antibody subtypes were detected using 100 μl/well Goat anti-human IgG1 (Mabtech, Cat: 3851-6-250), IgG2 (Mabtech, Cat: 3852-6-250), IgG3 (Mabtech, Cat: 38536-250) and IgG4 (Mabtech, Cat: 3854-6-250) biotinylated antibody subtypes (Mabtech, Sweden) seperately, diluted 1:1000 in PBS containing 1 % BSA. Plates were then incubated for 30 min at room temperature and Streptavidin-HRP (Mabtech, Cat: 3310-9) diluted to 1/1000 in PBS containing 0.1% BSA was added 100 μl/well for 30 min at room temperature.

TMB ELISA substrate solution (Mabtech, Cat: 3652-F10) was added 100 μl/well and incubated in the dark for 10 min at room temperature. The reaction was stopped by adding 100 μl/well 2M H_2_SO_4_ stop solution and absorbance values were read at 450nm.

### Statistical analysis

Statistical analysis was performed using Graph Pad Prism version 6. As the data were not normally distributed (as determined by the frequency distribution analysis of Graphpad PRISM), non-parametric tests were used in the statistical analysis. Differences in the serial values of NS1 antibodies, NS1 antigen and viral loads in patients with DHF and DF were calculated/compared using the Holm-Sidak method. Corrections for multiple comparisons were completed using the Holm-Sidak method and the statistical significant value was set at 0.05 (alpha). Differences in mean values between antibody responses to overlapping peptides in patients with DF and DHF and in individuals with SD and NSD were compared using the Mann-Whitney U test (two tailed).

**Table 1:** Clinical and laboratory characteristics of secondary dengue patients with DHF and DF recruited for the study.

| **Clinical findings** | **DHF (n= 33)** | **DF (n= 22)** |
| --- | --- | --- |
| Vomiting | 14 (42.42%) | 5 (22.72%) |
| Abdominal pain | 21 (63.63%) | 5 (22.72%) |
| Hepatomegaly | 28 (84.84%) | 0 |
| Bleeding manifestations | 1 (03.03%) | 3 (13.63%) |
| Pleural effusion | 18 (54.54%) | 0 |
| Ascites | 33 (100.0 %) | 0 |
| Lowest platelet count |  |  |
| <20,000 cells/mm<sup>3</sup> | 23 (69.69%) | 2 (9.09%) |
| 20,000 to 50,000 | 10 (30.30%) | 9 (40.90%) |
| 50,000-100,000 | 0 | 8 (36.36%) |
| >100,000 | 0 | 3 (13.63%) |
| Lowest Lymphocyte count |  |  |
| <750 | 21 (63.63%) | 9 (40.90%) |
| 750 – 1500 | 10 (30.30%) | 13 (59.09%) |
| >1500 | 2 (6.06%) | 0 |
| Infecting serotype |  |  |
| DENV1 | 10 (30.30%) | 10 (45.45%) |
| DENV2 | 23 (69.69%) | 12 (54.54%) |

## Data Availability

All data generated or analysed during this study is available included in the main manuscript and the supplementary files.

## Acknowledgements

Funding was provided by the Centre for Dengue Research, University of Sri Jayewardenapura, National Research Council, Sri Lanka (15-043), National Science Foundation, Sri Lanka (RPHS/2016/D-06) and by the Medical Research Council (UK). Graham Ogg receives support from the National Institute for Health Research (NIHR) Oxford Biomedical Research Centre (BRC).

## Author contributions

G.N.M and G.S.O designed the project and experiments, D.J and L.G performed the NS1 antibody experiments, S.F., L.G. and C.J. did the viral loads and other antibody experiments, C.J., P.G.S.B.J, D.H., M.A.P.A.P, S.F. and A.W recruited the patients and collected all clinical data, D.J, L.G and G.N.M analysed and interpreted the data, D.J., G.N.M, C.S.H. and G.S.O wrote the manuscript.

## Competing Interest

None.

